# Behavioral effects of developmental exposure to JWH-018 in wild type and disrupted in schizophrenia 1 (*disc1*) mutant zebrafish

**DOI:** 10.1101/2020.07.16.206615

**Authors:** Judit García-González, Bruno de Quadros, Alistair J. Brock, Caroline H. Brennan

## Abstract

Synthetic cannabinoids can cause acute adverse psychological effects, but the potential impact when exposure happens before birth is unknown. Use of synthetic cannabinoids during pregnancy may affect fetal brain development, and such effects could be moderated by the genetic makeup of an individual. Disrupted in schizophrenia 1 (*DISC1*) is a gene with important roles in neurodevelopment which has been associated with psychiatric disorders in pedigree analyses. Using zebrafish as a model, we investigated (1) the behavioral impact of developmental exposure to JWH-018 (a common psychoactive synthetic cannabinoid) and (2) whether *disc1* moderates the effects of JWH-018. As altered anxiety responses are seen in a several psychiatric disorders, we focused on zebrafish anxiety-like behavior. Zebrafish embryos were exposed to JWH-018 from one to six days post-fertilization. Anxiety-like behavior was assessed using forced light/dark and acoustic startle assays in larvae, and novel tank diving in adults. Compared to controls, developmentally exposed zebrafish larvae had impaired locomotion during the forced light/dark test, but anxiety levels and response to startle stimuli was unaltered. Adult zebrafish developmentally exposed to JWH-018 spent less time on the bottom of the tank, suggesting decreased anxiety. Loss-of-function in *disc1* increased anxiety but did not alter sensitivity to JWH-018. Results suggest developmental exposure to JWH-018 has behavioral impact in zebrafish, which is not moderated by *disc1*.

## 1. Introduction

In contrast to tobacco smoking, where prevalence during pregnancy has dropped from 14.6 to 10.6% in the United Kingdom [1], cannabis use among pregnant women has risen in recent years [2]. Cannabis does have medical utility for some conditions and may help pregnant women to alleviate nausea that usually accompanies pregnancy. However, cannabis may also affect fetal neurodevelopment, leading to long-term behavioral alterations [3]: The endocannabinoid system is present and plays an important role in early brain development [4]. Delta-9-tetrahydrocannabinol (THC) is the major psychoactive component of marijuana and can cross the placental barrier [3]. Thus, THC is able to bind the cannabinoid receptors located in the fetus brain, interfering with the endocannabinoid system and affecting neurogenesis and neuronal migration [3].

Similar to cannabis, synthetic cannabinoids commercialized as ‘Spice’, ‘K2’, ‘legal weed’ or ‘herbal incense’ gained popularity during the early 2000s and were legal in many countries for years [5]. The prevalence of synthetic cannabinoid consumption ranges between 0.2-4% in the general population [6], but prevalence estimates in pregnant women are unavailable, and it is likely that reported exposures are significantly underestimated.

JWH-018 (1-pentyl-3-(1-naphthoyl)indole) is one of the most common psychoactive synthetic cannabinoids. JWH-018 has high binding affinity for the cannabinoid receptors CB1 and CB2 [7,8] and mimics the physiological effects of THC through activation of the CB1 receptor [9]. Importantly, whereas THC is a partial agonist with weak affinity for CB1, JWH-018 is a full CB1/CB2 agonist with effects four to eight times more potent than THC [10,11]. Due to its potent effect, adverse outcomes associated with using synthetic cannabinoids containing JWH-018 may be more frequent and severe than those arising from cannabis consumption. Epidemiological studies show that acute intake of JWH-018 can cause strong psychological effects such as anxiety, psychosis, hallucination and alterations in cognitive abilities [12,13]. Given the potent adverse effects of acute exposure in adults, it is important to understand the short and long-lasting consequences of JWH-018 exposure during brain development. However, such consequences still remain unknown [14].

Genetic vulnerability to the effects of maternal drug intake during pregnancy may exacerbate adverse outcomes in the offspring. In particular, some genes that play important roles in neurodevelopment may modulate the effects of developmental exposure to drugs. Disrupted in Schizophrenia 1 (*DISC1*) is a gene in chromosome 1q42.1 that encodes a scaffolding protein with several protein interactions. Over 100 proteins have been suggested to interact with *DISC1* [15], highlighting the pivotal role of this protein during neurodevelopmental processes such as neuronal proliferation and migration, neuron spine formation, and synapse maintenance [15].

*DISC1* was identified in a Scottish family pedigree, where a translocation between chromosome 1 and 11 [(t(1;11)(q42.1;q14.3)] segregated with psychiatric disorders including schizophrenia, depression, and bipolar disorder [16,17]. The association between *DISC1* and psychiatric disorders was replicated in a second American pedigree with a 4 bp frameshift deletion in *DISC1* exon 12 [18]. However, there has been controversy regarding the relevance of this gene to psychiatric disorders as it seems likely that the association of *DISC1* with psychiatric disorders is driven by rare genetic variation that predisposes to psychiatric disorders only in certain individuals.

Despite the controversy about whether genetic variation in *DISC1* influences vulnerability to psychiatric disorders, there is consensus that *DISC1* plays an important role in neurodevelopment [15,19]. There is also some evidence suggesting that alterations due to *DISC1* loss-of-function are exacerbated by exposure to cannabinoids. *Disc1* mutant mice are more susceptible to deficits in fear-associated memory after exposure to THC during adolescence [20]. Perturbation of expression of *Disc1* in astrocytes, but not neurons, exacerbated the effects of adolescent THC exposure on recognition memory assessed in adult mice [21]. Altered expression of *Disc1* and THC exposure caused synergistic activation of the proinflammatory nuclear factor-k-B–cyclooxygenase-2 pathway in astrocytes, leading to secretion of glutamate and dysfunction of GABAergic neurons in the hippocampus [21]. These studies suggest that *Disc1* loss-of-function exacerbates the behavioral effects of THC exposure during adolescence, but no studies have yet examined the effects on earlier developmental exposures, nor the interaction of other cannabinoids (i.e. JWH-018) with *Disc1*.

Mammalian models such as rodents have been used to investigate early development and the effect of prenatal exposure to drugs of abuse (reviewed in [22]). Although these models are valuable, they present significant limitations: a mammalian fetus cannot be directly accessed and thus it is challenging to follow fetal neurodevelopment in vivo. In utero embryonic development makes it difficult to separate maternal and embryonic effects of exposure. Moreover, mammalian models are not suitable to fill the need for fast and high throughput screening of large numbers of compounds and mixtures, as well as multiple candidate biological pathways and their interactions. Using these models for experimental purposes would result in high costs of animal maintenance together with large space-requirements and relatively long gestation periods.

Zebrafish present important advantages over mammalian models [23]: firstly, embryos develop externally, and thus exposure is done directly through the water and not by maternal transfer. Secondly, embryonic development is only a five-day period from fertilization to a free-swimming and feeding larvae, therefore screening for potential neurobehavioral alterations is available within days of embryonic exposure. Thirdly, high fecundity of breeding adults provides sample sizes suitable for high-throughput screening experiments with multiple treatments/doses. Embryos/larvae fit into 96-well plates and are able to absorb small molecules through the skin, which removes issues regarding formulation. Furthermore, the embryos are transparent, which allows for easy monitoring of their development and for identifying abnormalities. Although zebrafish cannot develop human psychiatric disorders, they can display behaviors that resemble stress [24], anxiety [25] or drug seeking [26]. These behaviors are often called ‘intermediate phenotypes’ or ‘endophenotypes’ [27] and are assumed to be closer to the underlying genetic causes of psychiatric disorders [28]. Zebrafish are therefore an ideal animal model to investigate the short- and long-lasting effects of developmental exposure to drugs of abuse.

Our two main aims were to interrogate whether the developing central nervous system is susceptible to the effects of JWH-018, and to investigate whether loss-of-function mutations in the *disc1* gene exacerbates the effects of early developmental exposure to JWH-018. Using zebrafish as the animal model, we addressed the following research questions: (1) does developmental exposure to JWH-018 modulate behavior in larvae zebrafish?, (2) are the effects of developmental exposure to JWH-018 similar to the effects of THC and nicotine?, and (3) are the short- and long-lasting effects of developmental exposure to JWH-018 exacerbated by *disc1* loss of function?

## 2. Materials and Methods

### 2.1. Experimental design and timeline

Wild type zebrafish were exposed to 3 μM JWH-018 (Tocris, Cat. No. 1342), from 24 hours to six days post fertilization (dpf). At five dpf (with larvae being exposed to the drug for 96 hours), distances travelled during forced light/dark transitions were examined. Importantly, larvae were *in* the drug solution during behavioral testing, and drug was refreshed 3-5 hours prior to placing the animals into the Danio Vision Observation Chamber. At six dpf (with larvae being exposed to the drug for 120 hours), response and habituation to acoustic startle stimuli were examined. Larvae were also *in* the drug solution during the response and habituation to startle stimuli test, but in this case the drug solution was not refreshed prior to testing.

To investigate whether the effect of JWH-018 was similar to other psychoactive substances with well characterized effects on zebrafish (namely THC and nicotine), we repeated the experimental protocol and behavioral battery in wild type zebrafish larvae using 2 μM THC (Merck, Cat. No. T4764), and 0.15 μM nicotine (Sigma, Cat. No. N1019). Drugs were refreshed with the same time course.

To examine the potential interactions between JWH-018 exposure and *disc1* mutations in the short and long term, we repeated the developmental exposure to 3 μM JWH-018 using *disc1* wild type and mutant zebrafish and their behavior was assessed at five and six dpf (as in experiments with wild type zebrafish). Furthermore, *disc1* wild type and mutant zebrafish treated with JWH-018 but not used for larval behavioral testing were reared to adulthood in normal conditions. At four months old, the anxiety-like response of the exposed vs non-exposed fish was assessed using the novel tank diving procedure. An overview of the study design and experimental timeline is represented in Figure 1.

**Figure 1.**
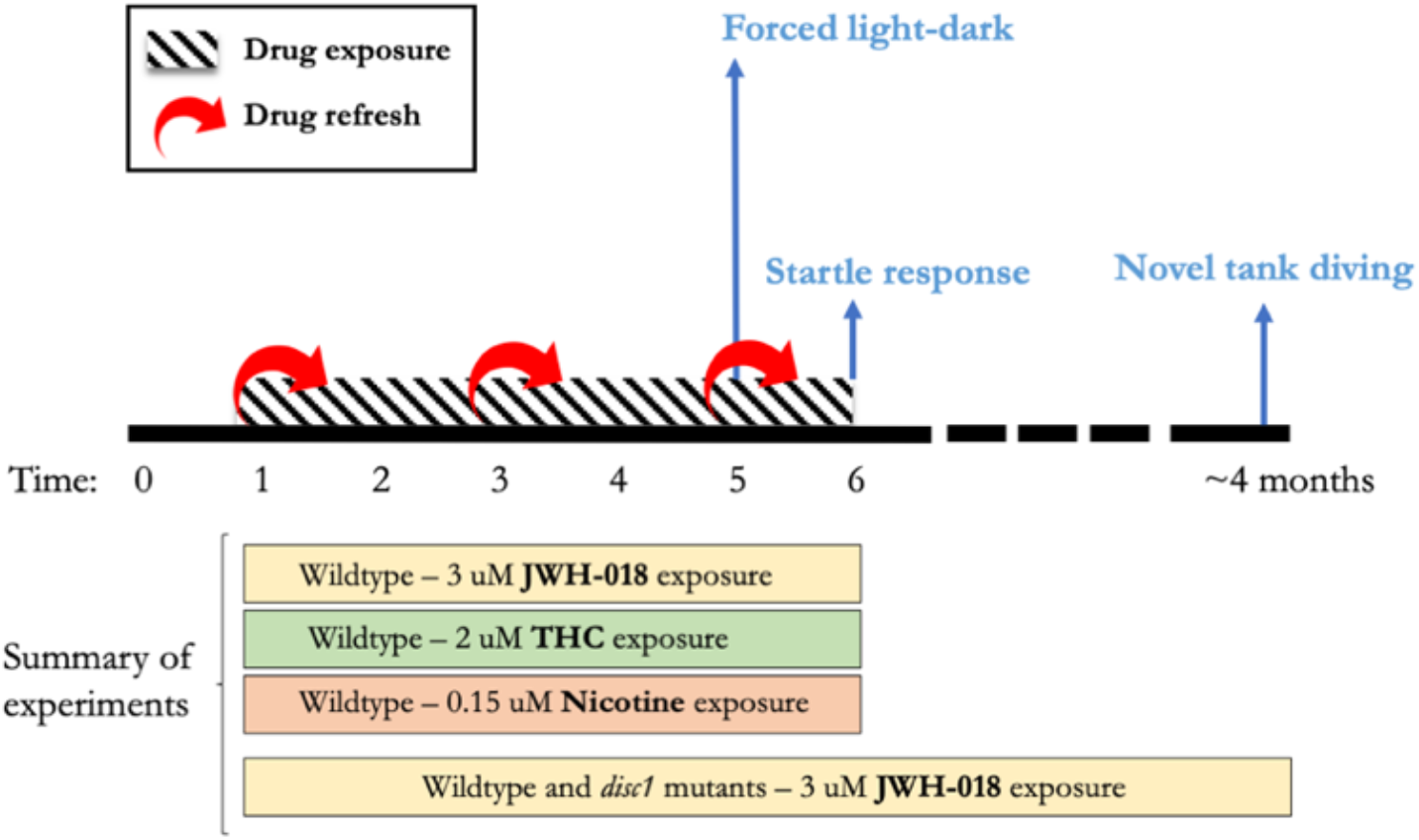
Experimental timeline for developmental exposure to JWH-018, THC, and nicotine. Horizontal bars in the lower part of the figure represent experiments carried out. The behavioral tests performed are represented in light blue.

### 2.2. Animal maintenance

Zebrafish were housed in a recirculating system (Techniplast, UK) on a 14hour:10hour light:dark cycle (08:30–22:30). The housing and testing rooms were at ~25–28°C. Zebrafish were maintained in aquarium-treated water and fed three times daily with live artemia (twice) and flake food (once). Wild type zebrafish belonged to the Tübingen strain. The *disc1* line (AB background strain) was obtained from the Cecilia Moens lab (Fred Hutchinson Cancer Research Center, Seattle, USA), and was provided by Dr Jon Wood (University of Sheffield). The mutant allele (*disc1;^fh291^*) is caused by a point mutation in exon 2 (T>A), that produces an early stop codon. More information is detailed elsewhere [29].

To breed zebrafish, we placed them in breeding tanks which had either perforated floors or a container with marbles to isolate eggs from progenitors. We moved the animals to breeding tanks in the evening and collected eggs the following morning. Eggs were incubated in Petri dishes at 28°C until five dpf. If reared, larvae were moved to the recirculating system at six dpf and fed with commercial fry food.

All procedures were carried out under license in accordance with the Animals (Scientific Procedures) Act, 1986 and under guidance from the local animal welfare and ethical review board at Queen Mary University of London.

### 2.3. Developmental drug exposure

#### 2.3.1. Developmental exposure to JWH-018, THC and nicotine in wild type Tübingen larvae

Since JWH-018 and THC are not soluble in water, JWH-018 was dissolved in DMSO (Sigma-Aldrich, Cat. No. D8418), and THC was provided by the manufacturer in methanol (MeOH). Care was taken to ensure that the final carrier concentration for all samples was 0.1% DMSO (for JWH-018 experiments) and 0.01% MeOH (for THC experiments). To account for potential effects of the carrier substance, we used 0.1% DMSO and 0.01% MeOH respectively as control groups. Drug and control solutions were changed every 48 hours to ensure constant drug uptake by the zebrafish embryos and to account for oxidation in the water.

Drug concentrations for JWH-018 and THC were chosen based on previous studies, where exposure to 2 μM THC led to impaired locomotor response in zebrafish larvae [30], and 3 μM JWH-018 led to behavioral alterations in rodents [31,32]. Developmental exposure to 0.15 μM nicotine was chosen because previous studies in our lab showed this dose induced increased nicotine preference in adult zebrafish (Appendix A and supplementary Figure 3).

#### 2.3.2. Developmental exposure to JWH-018 in *disc1* mutant larvae

Exposure to 3 μM JWH-018 and behavioral testing at five and six dpf using *disc1* wild type and mutant zebrafish was carried out as for the wild type larvae. Larvae were obtained from an in cross of *disc1* heterozygous zebrafish. Therefore, larvae were a mix of wild type, homozygous and heterozygous zebrafish that were randomly allocated in the experimental plates and genotyped after behavioral testing. We performed five independent experiments on five different days. To account for variation across experiments/days, the date of testing was included as a covariate in the analyses.

### 2.4. Behavioral assays

#### 2.4.1. Forced light/dark test

The forced light/dark test is a well-established behavioral assay in zebrafish larvae, where changes in locomotor activity due to alternating bright light/dark depend on the integrity of brain function and the correct development of the visual and nervous system. Transitions from dark to bright light cause an abrupt decrease in larval movement (freezing), and the subsequent progressive increase in movement can be interpreted as a measure of recovery to stress-reactivity and anxiety [33].

We conducted forced light/dark tests between 9 am and 4 pm with the drug present in the water. We placed larvae in 48-well plates. To reduce stress due to manipulation, we let them acclimate for at least one hour in ambient light before testing. Larvae were exposed to alternating light dark cycles of 10 min: there was an initial 10 minutes period of dark (baseline), followed by two cycles of 10 minutes of light and 10 minutes of dark. This protocol has been used elsewhere [34]. Distances travelled were recorded using Ethovision XT software (Noldus Information Technology, Wageningen, NL) and data were outputted in one-minute time-bins. Data was fitted to linear mixed models with total distance travelled as response variable, experimental variables (e.g. genotype, dose, time) as fixed effects, and fish ID as random effects. Details on the data analysis is detailed in Appendix B.

#### 2.4.2. Response and habituation to startle stimuli test

In response to abrupt sound/vibration stimuli zebrafish larvae execute a fast, non-associative learning escape response. This response has been extensively characterized and involves one of two distinct motor behaviors: a short-latency C-bend of the tail, initiating within 5–15 milliseconds of the stimulus, or a slower, long-latency C-bend response initiating within 20–80 milliseconds. These two motor behaviors use different, possibly overlapping neuronal circuitry [35] but in this study they were measured jointly, since a high-speed camera was not available.

When the abrupt sound/vibration stimuli are given repeatedly, zebrafish exhibit iterative reduction in the magnitude of the response, commonly known as habituation. Habituation is the mechanism by which the nervous system filters irrelevant stimuli. It is evolutionarily conserved and present in a wide range of species from invertebrates, such as Aplysia and Drosophila, to vertebrates such as rodents [36]. Defective habituation is also associated with neuropsychiatric disorders such as schizophrenia [37].

We assessed the response and habituation to startle stimuli between 9 am and 4 pm with the drug present in the water (but without drug refresh prior to the test). We used the DanioVision Observation Chamber, which contains a dedicated tapping device, and set the DanioVision tap stimulus at the highest intensity (intensity level: 8). Larvae were subjected to 10 sound/vibration stimuli over 10 seconds (1 second interval between each stimulus). For all experiments, distance travelled was recorded using Ethovision XT software (Noldus Information Technology, Wageningen, NL) and data were outputted in one second time-bins.

As proof of concept, we replicated the experiment by Best and colleagues [38], where 50 stimuli were given using 1, 5 and 20 seconds inter-stimulus intervals (ISI). Following the habituation paradigm [36], shorter ISI led to faster habituation [Effect of ISI: χ2(2)=19.04, p<0.0001] (Figure S1).

#### 2.4.3. Novel tank diving test

Novel tank diving exploits the natural tendency of zebrafish to initially stay at the bottom of a novel tank, and gradually move to upper parts of the tank. The degree of ‘bottom dwelling’ has been interpreted as an index of anxiety (greater bottom dwelling meaning greater anxiety) and it is conceptually similar to the rodent open-field and elevated plus maze tasks [25]. Other measures such as the distance travelled in the tank during the course of the assay and the transitions to bottom of the tank can give further insights on the hyper-responsiveness to novel environments.

We transported adult zebrafish (3-4 months) to the behavioral room in their housing tanks and let them acclimate to the room conditions for at least one hour before testing. Novel tank diving was assessed as previously described [39]: zebrafish were individually introduced into a 1.5 L trapezoid tank (15.2 cm x 27.9 cm x 22.5 cm x 7.1 cm) (Figure S2) and filmed for five minutes. Their behavior was tracked using EthoVision system (Noldus, Netherlands) and data were outputted in one-minute time-bins. Care was taken to ensure that experimental groups were randomized during testing. Behavioral testing was conducted between 9 am and 2 pm.

We analyzed three behaviors in response to the novel tank: (1) time that zebrafish spent on the bottom third of the tank, (2) total distance that zebrafish travelled in the tank over the five minutes, and (3) number of transitions to the top-bottom area of the tank. Details on the data analysis are in Appendix B.

#### 2.4.4. Code availability

Code used to analyze the behavioral assays is available at https://github.com/juditperala/Zebrafish-behaviour.

### 2.5. Competitive allele-specific PCR (KASP^™^) disc1 larvae genotyping

After behavioral testing, DNA was extracted using the hot shock DNA extraction protocol. Since the loss-of-function in *disc1* is caused by a point mutation, we used the competitive allele-specific PCR (KASP™) assay (LGC, Biosearch Technologies) to genotype the zebrafish.

**Table 1.**
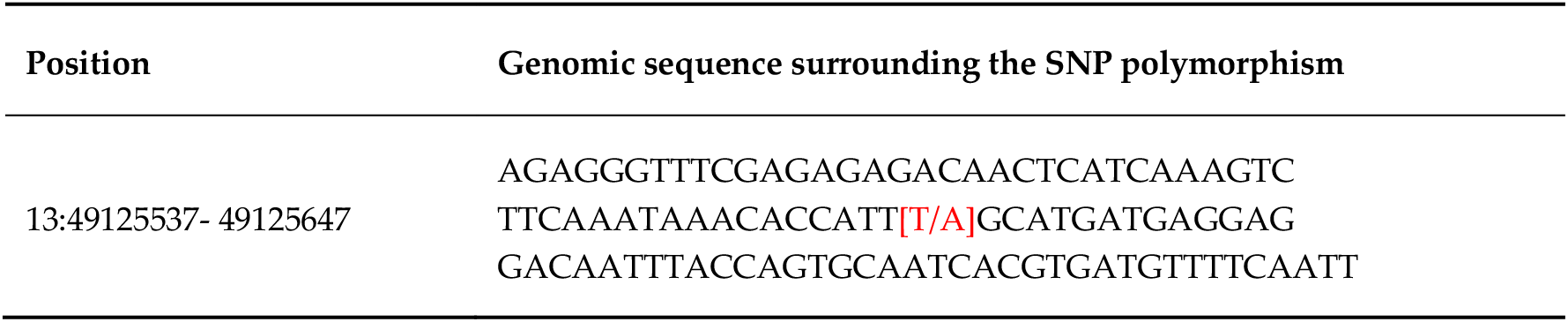
Genomic sequence surrounding the point loss-of-function mutation (T >A, in red) for *disc1*.

## 3. Results

### 3.1 Effects of developmental exposure to JWH-018 on larval behavior

#### 3.1.1. Forced light/dark test

Over the course of the forced light/dark test, time [χ2(1)=41.27, p<0.0001] and JWH-018 treatment [χ2(1)=17.53, p<0.0001] predicted distance travelled by five dpf larvae (Figure 2). Exposure to 3 μM JWH-018 impaired locomotion during baseline and dark periods. During the first minutes of the experiment, treated larvae travelled shorter distances (M=0.40, SE=0.04) than controls (M=0.50, SE=0.40) [*Effect of treatment during baseline*: χ2(1)=0.04, p=0.04]. Over the course of the two dark periods, control larvae sharply increased their locomotion and progressively reduced it, whereas larvae treated with 3 μM JWH-018 did not show as great an increase in movement (M=0.32, SE=0.03) as controls (M=0.55, SE=0.03) [*Effect of treatment during Dark1 and Dark2*: χ2(1)=30.88, p<0.0001].

**Figure 2.**
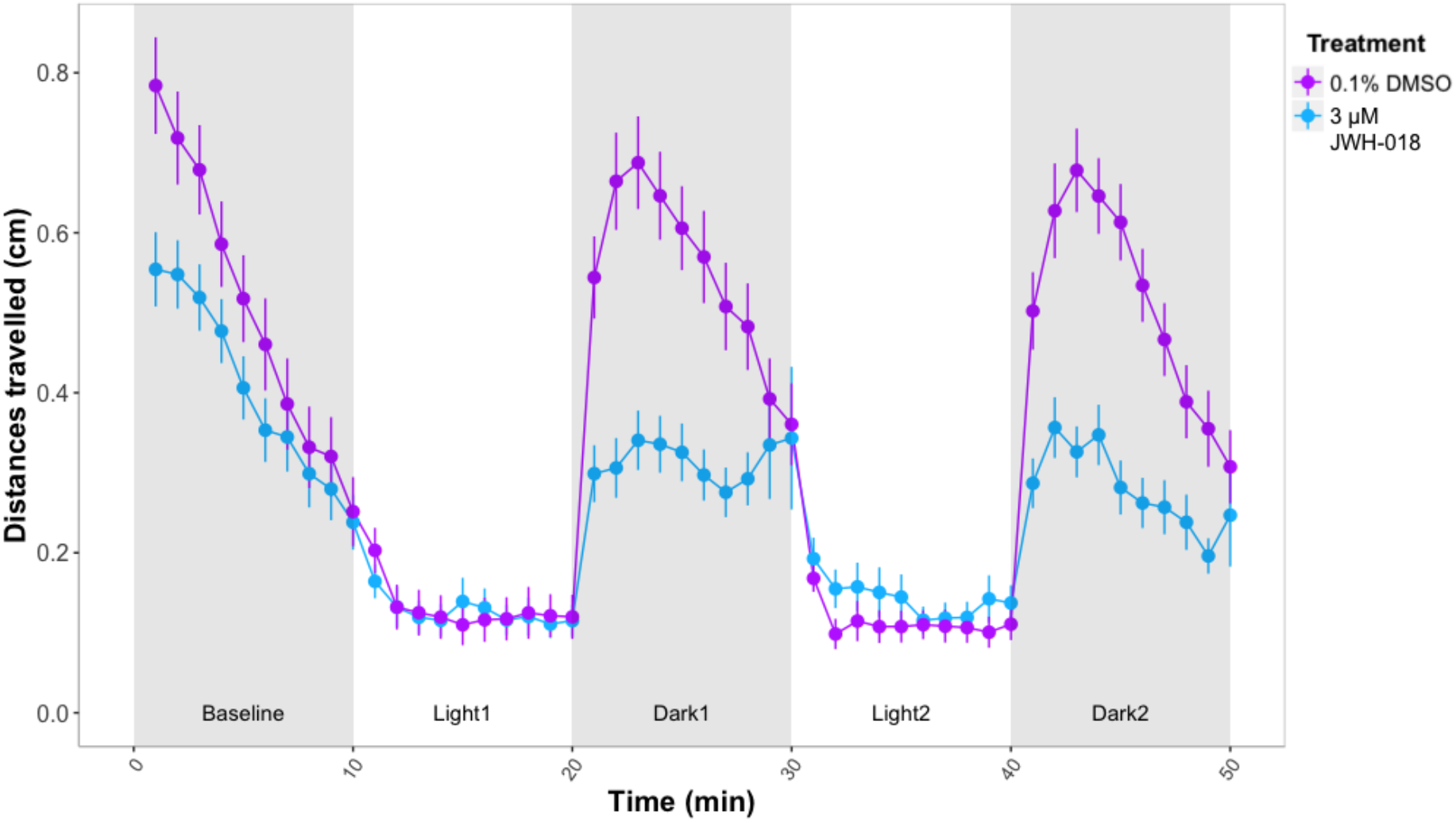
Forced light/dark test in five dpf zebrafish larvae. Sample size: n=64 for each dose group. Each dot represents mean distance travelled per minute. Error bars represent ±SEM.

The increase in locomotion during the light periods (measured as the slopes from minute 10 to 20 for the first light period, and minute 30 to 40 for the second light period) were interpreted as a measure of recovery to stressful stimuli and anxiety-like behavior. No significant differences between the slopes of treated vs control larvae were observed for any of the two light periods (p>0.05).

#### 3.1.2. Response to repeated sound/vibration startle stimuli

We next assessed the response to repeated startle stimuli at six dpf. There were no significant differences between 3 μM JWH-018 treated and control larvae in distance travelled before and during the stimuli (p>0.05) (Figure 3).

**Figure 3.**
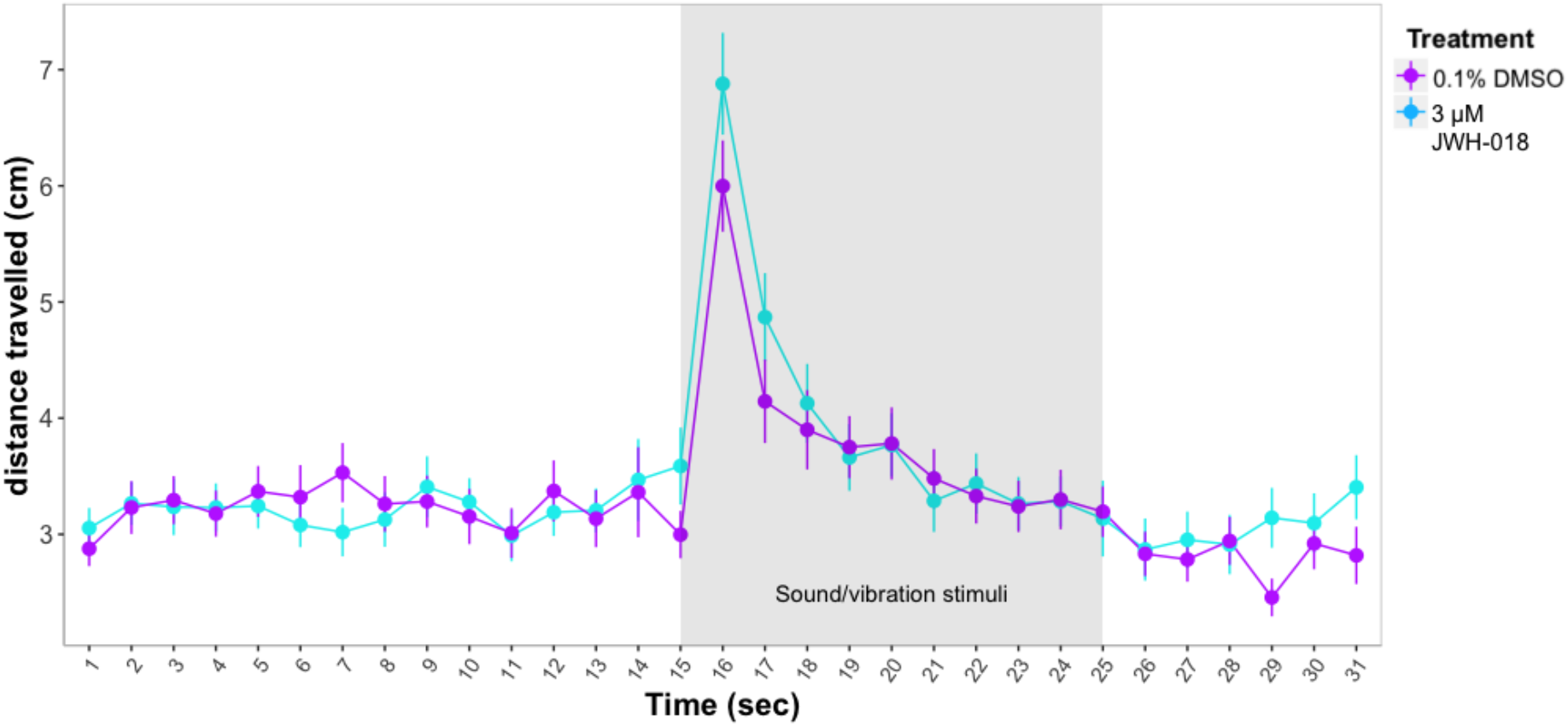
Response and habituation to startle stimuli test in six dpf zebrafish larvae. Sample sizes: control: n=87, JWH-018 treated: n=81. Each dot represents mean distance travelled per second. Error bars represent ±SEM.

### 3.2. Effects of developmental exposure to THC and nicotine on larval behavior

#### 3.2.1. Forced light/dark test

We investigated whether the behavioral effects of developmental exposure to nicotine and THC where similar to those of JWH-018. Exposure to 2 μM THC led to impaired locomotion of larvae, similar to the effects observed for the JWH-018 treatment. Distances travelled over the course of the experiment were much shorter for THC treated larvae (M=0.62, SE=0.02) compared to controls (M=0.91, SE=0.02) [*Effect of THC treatment*: *χ*2(1)=120.89, p<0.0001]. The differences between treated vs control larvae were consistent for baseline, light and dark periods (Figure 4).

**Figure 4.**
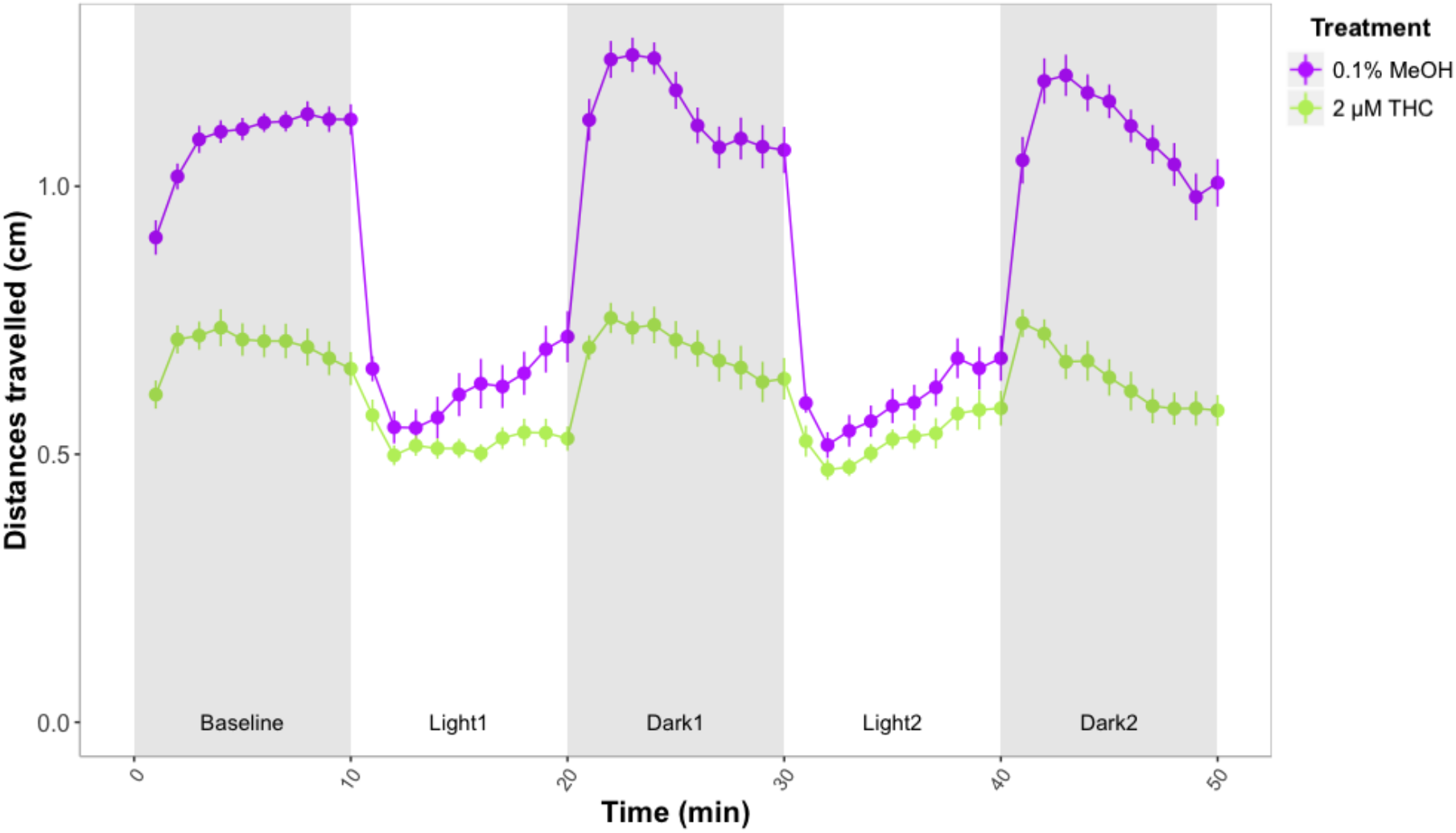
Forced light/dark test in wild type zebrafish exposed to 2 μM THC. Sample size: n=48 for each dose group. Each dot represents mean of the total distance travelled per minute. Error bars represent ±SEM.

Treatment with 2 μM THC also affected larvae recovery slopes during the first light period. Slopes for control larvae were steeper (M=0.02, SE=0.006) than for THC treated larvae (M=0.004, SE=0.006), suggesting that controls recovered faster and therefore THC may have an anxiogenic effect [F(1)=5.397, p=0.0223]. However, there were no significant differences between slopes of treated vs control larvae for the second light period (p>0.05).

In contrast to JWH-018 and THC, exposure to nicotine produced an increase in distances travelled. During the forced light/dark test, both time [*χ*2 (1)=15.56, p<0.0001] and nicotine treatment [χ2 (1)=16.04, p<0.0001] had a significant effect on the distance travelled over the course of the forced light/dark assay. Treatment with 0.15 μM nicotine increased the locomotor activity of larvae. The increased distances travelled by nicotine-treated larvae were significant for baseline, dark and light periods (p<0.0001) (Figure 5).

**Figure 5.**
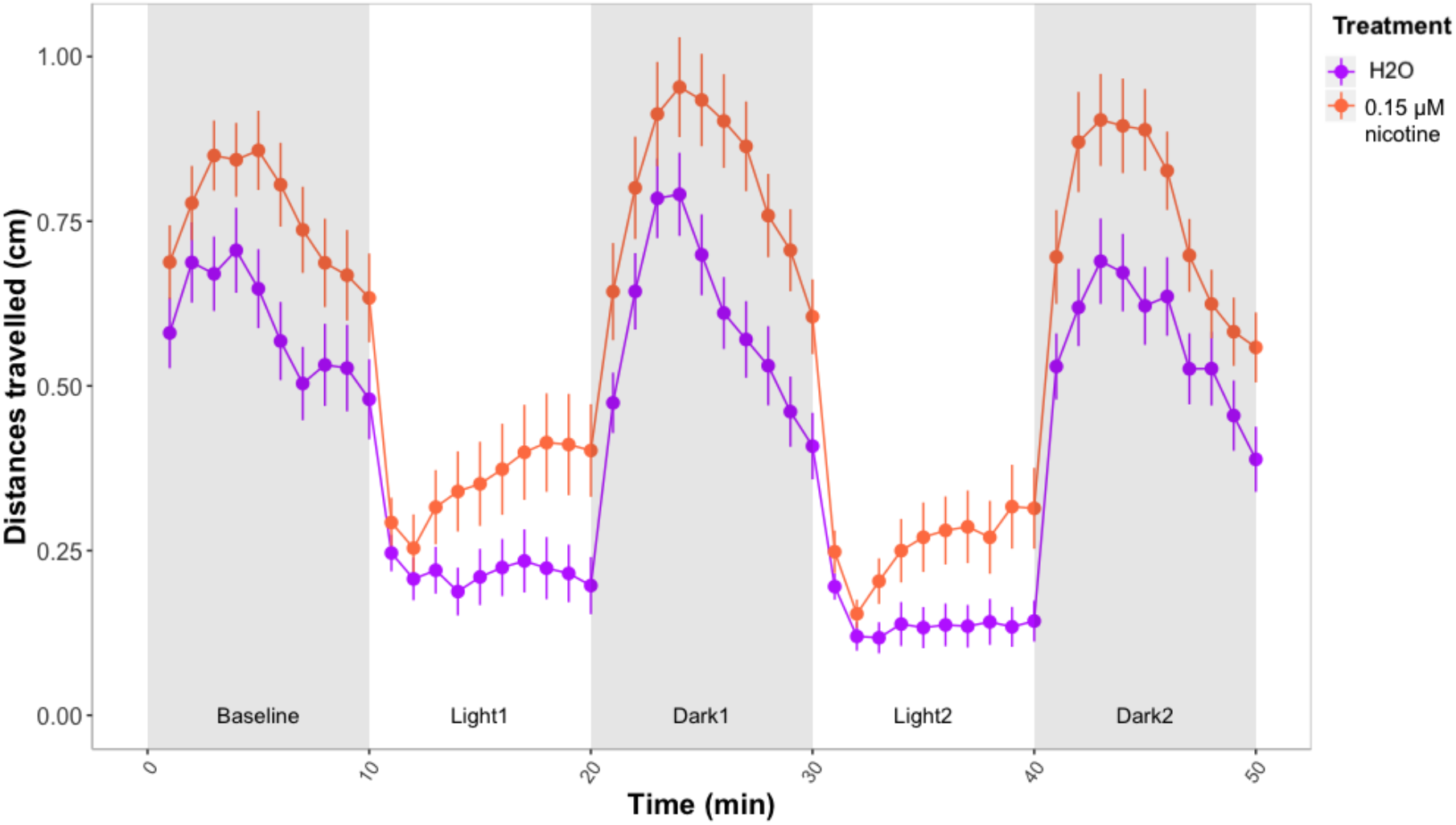
Forced light/dark test in wild type zebrafish exposed to 0.15 μM nicotine. Sample size: n=48 for each dose group. Each dot represents mean of the total distance travelled per minute. Error bars represent ±SEM.

There was a qualitative difference between control and treated zebrafish in the slopes during light periods, as nicotine-treated zebrafish seemed to recover faster, suggesting an anxiolytic effect of nicotine. However the difference between nicotine treated and control zebrafish was not significant [F(1)=3.18, p=0.07].

#### 3.2.2. Response to repeated sound/vibration stimuli

Zebrafish larvae treated with 2 μM THC were less active during the first 30 seconds of the experiment, before any stimuli was given [*Effect of THC treatment*: *χ*2(1)=15.31, p<0.0001]. However, during the ten sound/vibration stimuli larvae had similar locomotor activity (p>0.05) (Figure 6).

**Figure 6.**
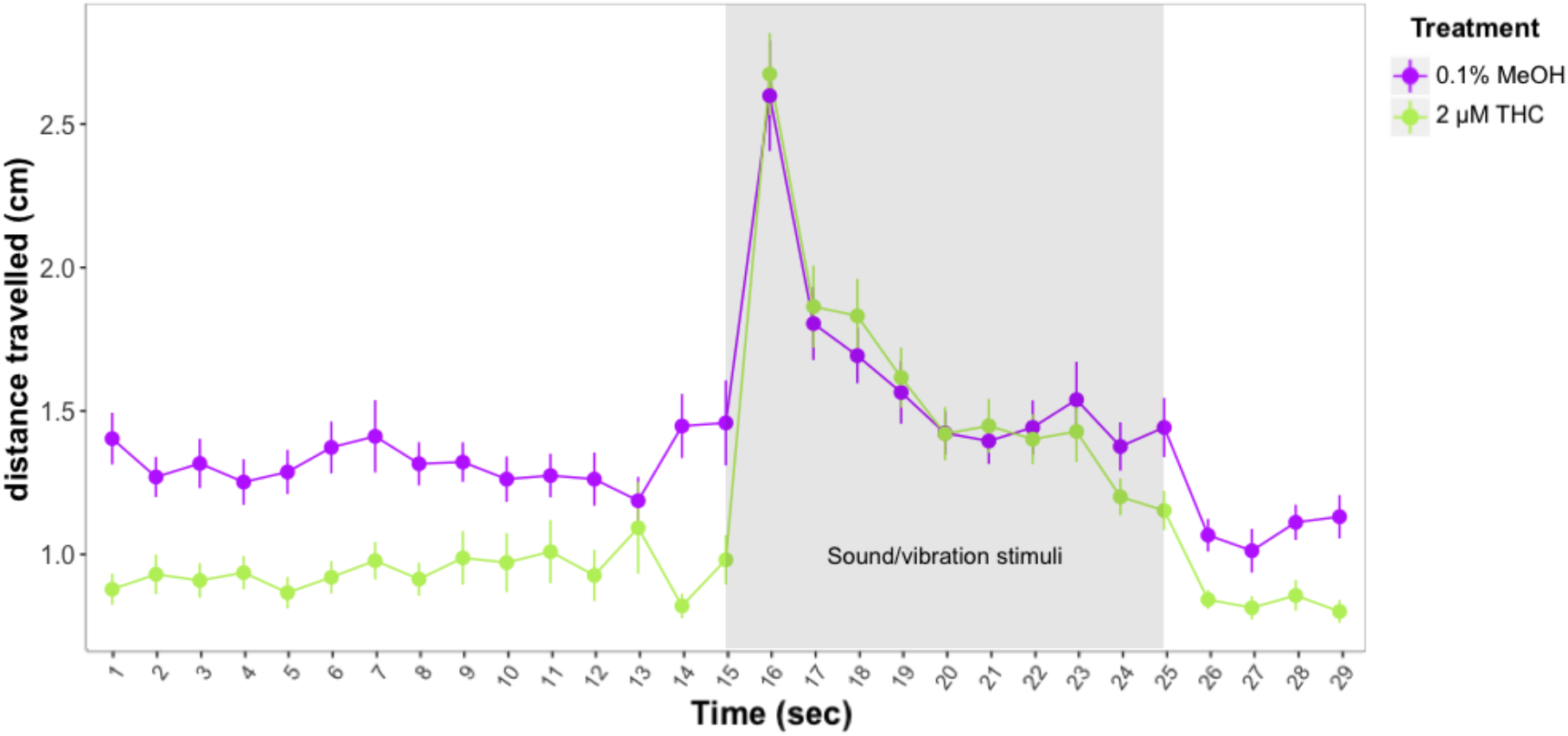
Distances travelled by control and THC treated larvae before and after exposure to 10 sound/vibration stimuli. Figure shows mean distances travelled in one second time bins. Error bars represent ±SEM. Sample sizes: n=48 per dose group.

Similar to the response seen during the forced light/dark test, zebrafish treated with 0.15 μM nicotine increased their locomotor response. The effect was significant during stimuli [*χ*2(1)=4.00, p=0.04], but not during the first 15 seconds before the stimuli (p>0.05) (Figure 7).

**Figure 7.**
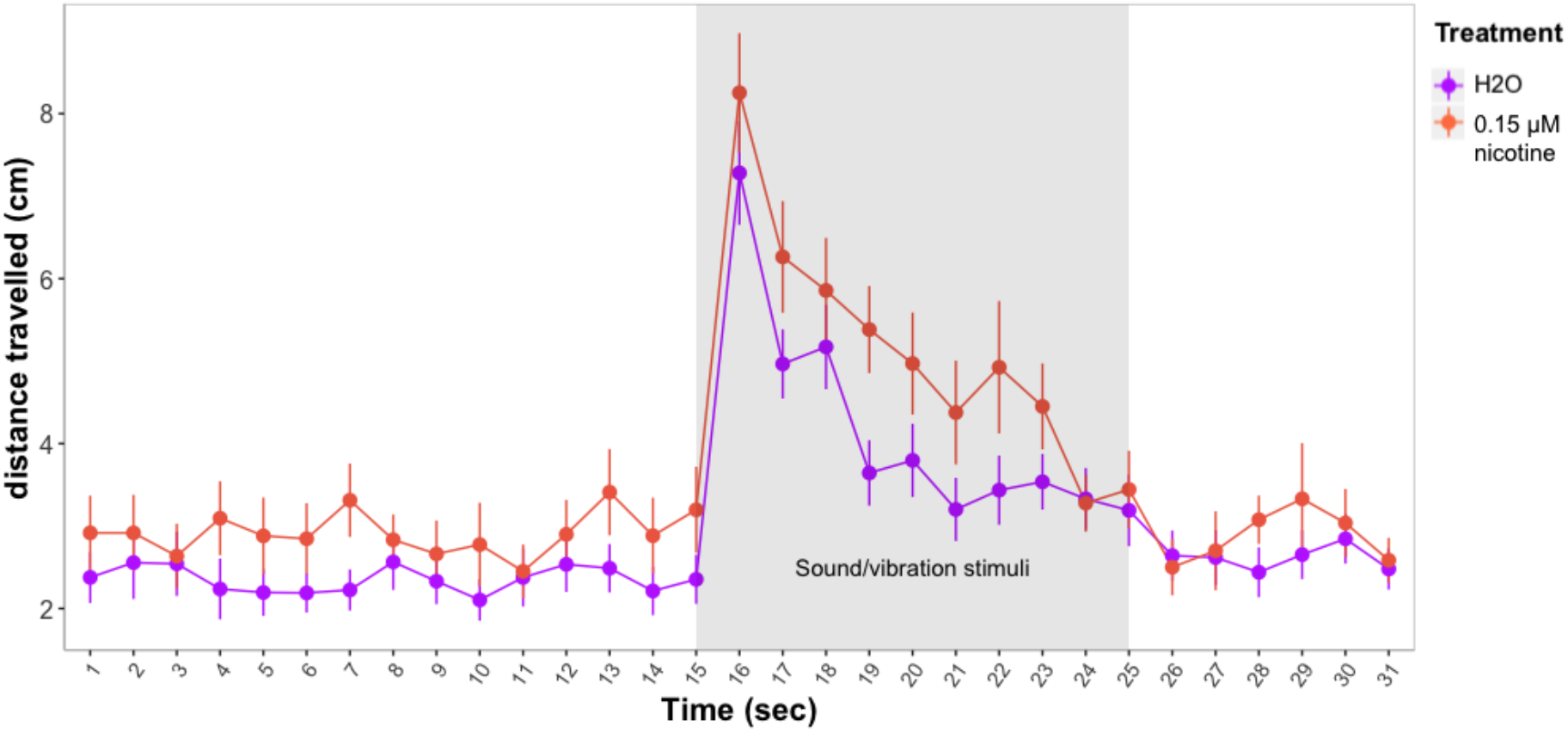
Distances travelled by control and nicotine treated larvae before and after exposure to 10 sound/vibration stimuli. Figure shows mean distances travelled in one second time bins. Error bars represent ±SEM. Control: n=23, treated with 0.15 μM nicotine: n=23.

### 3.3. Larval behavior during developmental exposure to JWH-018 in wild type and mutant disc1 larvae

Similar to the results for the Tübingen larvae, over the 50 minutes of the forced light/dark test, JWH-018 treatment [*χ*2(1)=12.51, p<0.0001] and time [*χ*2(1)=72.83, p<0.0001] were significant predictors of distance travelled. Although *disc1* wild type larvae travelled longer distances than mutants, genotype effects were not significant [χ2(1)=4.9, p=0.08] (Figure 8).

**Figure 8.**
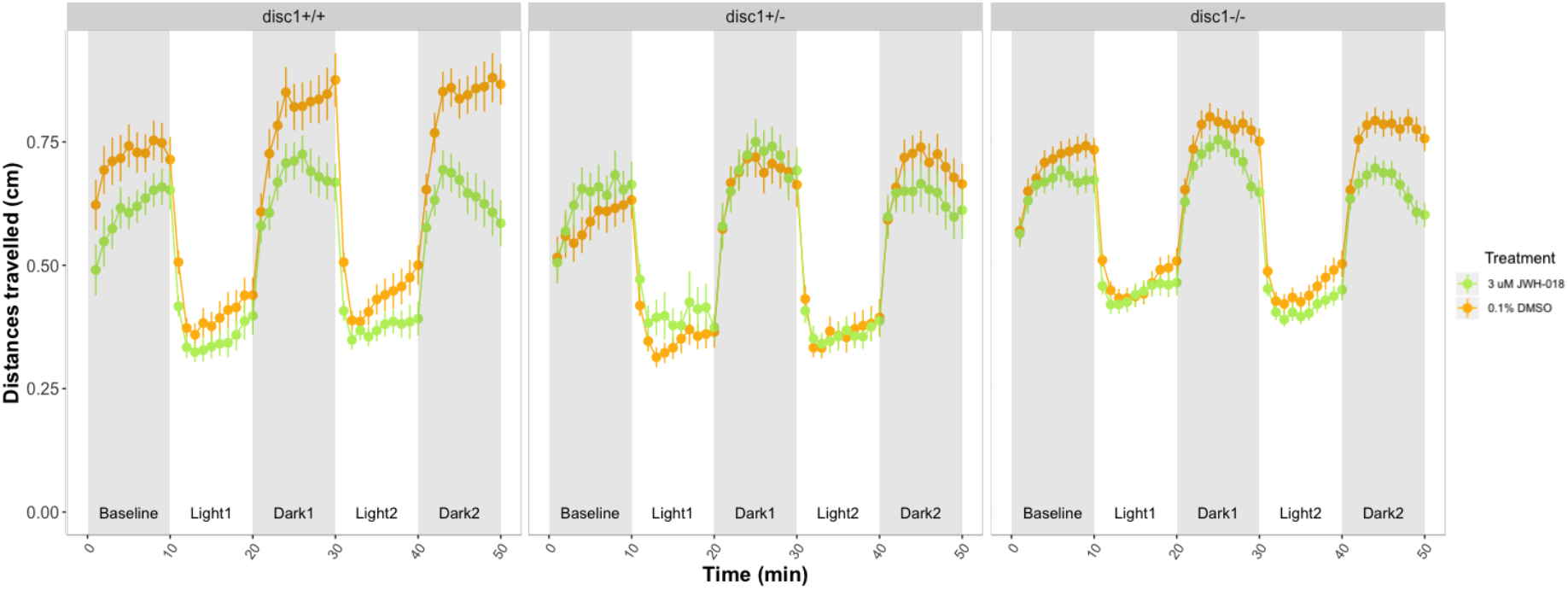
Forced light/dark test in five dpf wild type and *disc1* loss-function mutant larvae. Sample sizes for each group: control *disc1* +/+: n=30, JWH-018 *disc1* +/+: n=34, control *disc1* +/-: n=33, JWH-018 *disc1* +/-: n=27, control *disc1* -/-: n=107, JWH-018 disc1 -/-: n=92. Each dot represents mean distance travelled per minute. Error bars represent ±SEM.

During baseline, neither treatment nor genotype affected distances travelled (p>0.05). During the dark periods, wild type and *disc1* homozygous (but not *disc1* heterozygous larvae) travelled shorter distances when exposed to JWH-018 [*Effect of JWH-018 treatment*: *χ*2(1)=16.17, p<0.0001].

During light periods, there was a main effect of JWH-018 treatment [*χ*2(1)=4.57, p=0.032]: larvae exposed to JWH-018 travelled shorter distances than control larvae. However, there were no significant main effects of *disc1* genotype, nor significant interactions between genotype and JWH-018 on distances travelled or on th slopes calculated during light periods.

After 24 hours from the last JWH-018 drug refresh, treated and control larvae showed no significant differences in distances travelled before or during the startle stimuli. There were no significant differences across *disc1* genotype groups (Figure 9).

**Figure 9.**
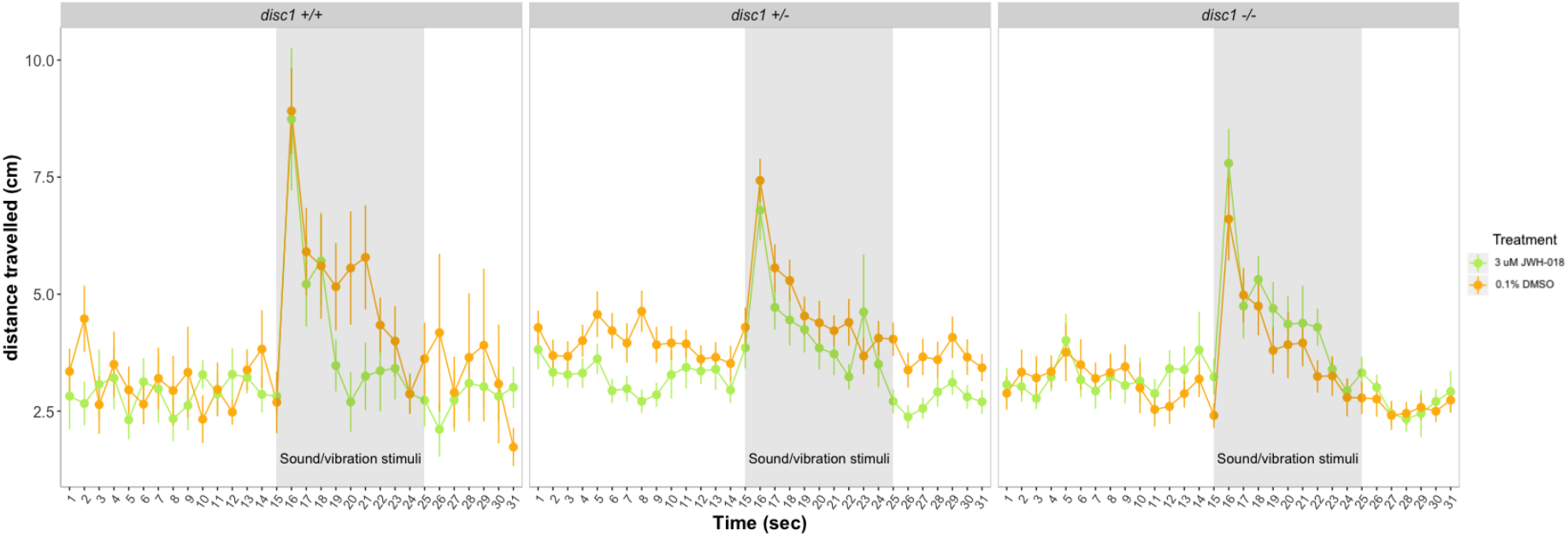
Response and habituation to startle stimuli test in six dpf control and JWH-018 treated wild type and *disc1* mutant larvae. Sample sizes: control *disc1* +/+: n=15, JWH-018 *disc1* +/+: n=13, control *disc1* +/-: n=47, JWH-018 *disc1* +/-: n=47, control *disc1* -/-: n=22, JWH-018 *disc1* -/-: n=22.

### 3.4. Adult behavior after developmental exposure to JWH-018 in wild type and mutant disc1 zebrafish

The *disc1* genotype affected the behavioural response during the novel tank assay (Figure 10). Wild type zebrafish spent less time on the bottom of the tank than homozygous and heterozygous *disc1* mutants [*Effect of genotype*: *χ*2(14)=119.40, p<0.0001] (Figure 10-A). Distances travelled over the five minutes of the experiment were also different across *disc1* genotypes (Figure 10-B): while wild type zebrafish did not differ in the distance travelled over time, zebrafish heterozygous and homozygous for *disc1* moved less during the first minute, and increased later the distance travelled [*Effect of genotype by time interaction*: χ2(14)=18.15, p=0.02]. The number of transitions between the bottom and top area of the tank over the five minutes of the experiment remained similar for wild types but increased for heterozygous and homozygous zebrafish *[Effect of genotype by time interaction:* χ2 (8)=22.93, p <0.0001] (Figure 10-C).

**Figure 10.**
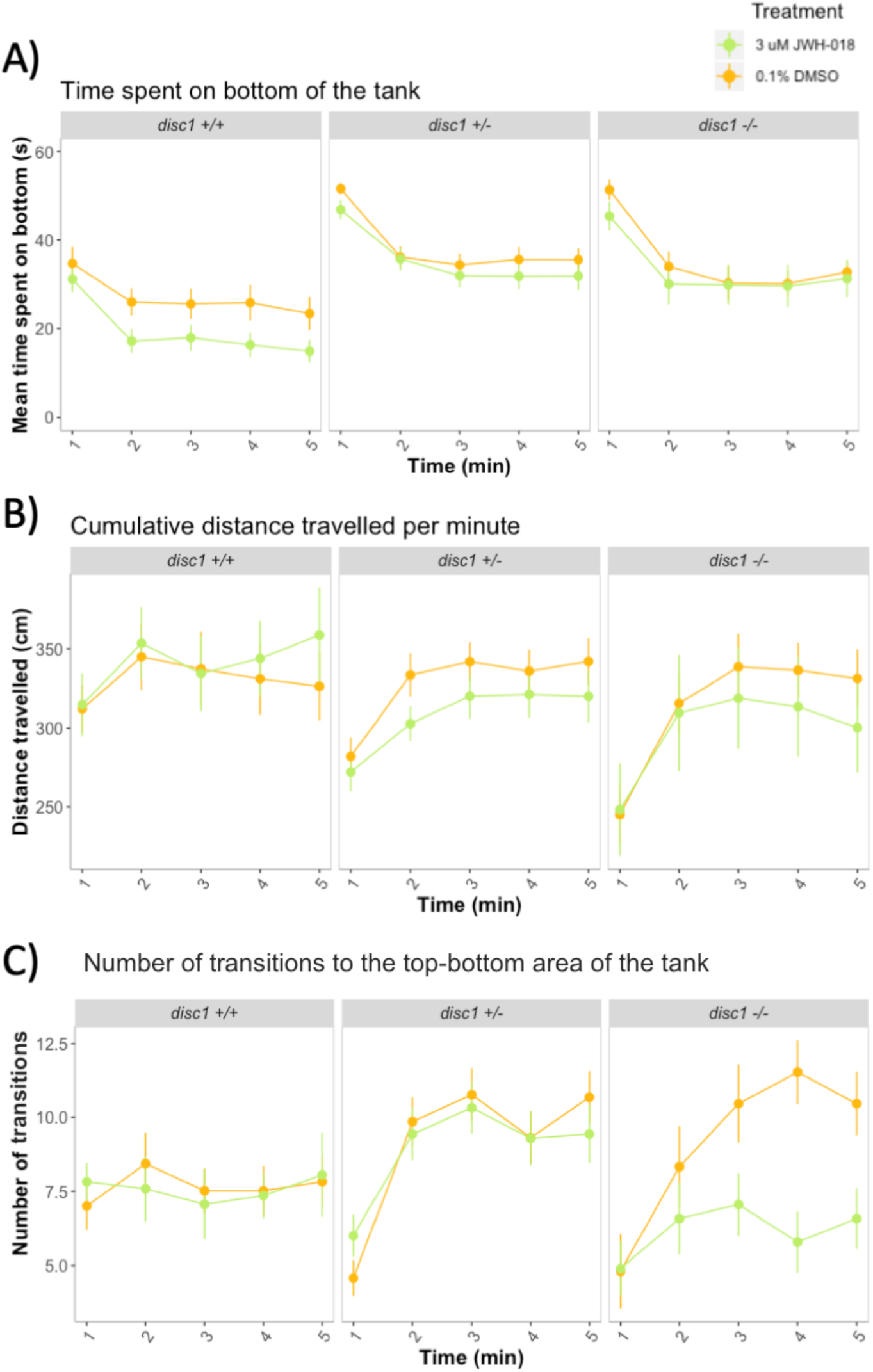
Novel tank diving response in adult wild type and mutant *disc1* zebrafish after developmental exposure to 3μM JWH-018. Sample sizes for each group: control *disc1* +/+: n=23, JWH-018 *disc1* +/+: n=17, control *disc1* +/-: n=35, JWH-018 *disc1* +/-: n=34, control *disc1* -/-: n=15, JWH-018 *disc1* -/-: n=19. Error bars represent ±SEM.

Developmental exposure to JWH-018 reduced the time spent on the bottom of the tank [*Effect of JWH-018 treatment*: χ2(1)=11.31, p<0.0001]. The effect was stronger for wild type than for mutant zebrafish (Figure 10-A), but there were no significant genotype by JWH-018 treatment interactions (p>0.05). Developmental exposure to JWH-018 did not affect the distance travelled nor the number of transitions between the top and bottom area of the tank for wild type and heterozygous *disc1* zebrafish (Figure 10-B and C) (p>0.05). For homozygous *disc1* zebrafish, treatment with JWH-018 decreased the number of top-bottom transitions but the interaction between genotype and JWH-018 treatment was not significant (Figure 10-C).

## 4. Discussion

This study used zebrafish as an animal model to investigate the behavioral effects of developmental exposure to JWH-018, the main psychoactive compound of synthetic cannabinoids. Zebrafish larvae exposed to JWH-18 had impaired locomotor response during the forced light/dark test but their anxiety levels and response to repeated sound/vibration stimuli were not altered. We then interrogated whether the behavioral effects of developmental exposure to JWH-018 were exacerbated by loss-of-function mutations in *disc1*, an important gene for neurodevelopment with a potential role in the cannabinoid system. Loss-of-function in *disc1* increased zebrafish anxiety but did not moderate sensitivity to the effects of JWH-018.

Alterations in the typical response to light and dark periods can be used to study the anxiety-like response in zebrafish. Others have interpreted the distance travelled in the dark period immediately following light exposure as a measure of anxiety -the greater the distance moved, the more anxious [40]. However, this interpretation is usually applied when using shorter light exposures (50 seconds) and is problematic when there are clear effects on locomotion. In this study, we examined the slopes during light periods, which represent how quickly zebrafish larvae recover from a startle stimulus (i.e. bright light) and provide a measure of stress and anxiety less biased by locomotor effects. Our results show developmental exposure to JWH-018 did not affect the recovery during light, suggesting no effects of JWH-018 on anxiety. By contrast, larvae exposed to THC recovered slower -suggesting an anxiogenic effect of THC, and larvae exposed to nicotine tended to recover faster -suggesting an anxiolytic effect of nicotine. Both THC and nicotine were used as positive controls, and our results are consistent with previous studies of the novel tank diving response in adult zebrafish: compared to controls, animals pre-exposed to THC spent more time on the bottom of the tank, consistent with an anxiogenic effect [41], whereas animals pre-exposed to nicotine spent less time on the bottom of the tank, consistent with an anxiolytic effect [42]. Since both THC and JWH-018 are cannabinoids, the difference in their behavioral impact is of interest. Differences in pharmacological properties (JWH-018 is a full CB1/CB2 agonist, whereas THC is a CB1 partial agonist) or pharmacokinetic warrant further investigation.

In addition to the anxiety-like behaviors, the stimulatory and depressant responses elicited by neuroactive drugs used by humans can be modeled in zebrafish larvae. For example, exposure to adrenaline -a neuro-stimulant-increased the locomotor activity in the forced light/dark test, whereas tricaine -a CNS depressant-decreased it [43]. In this study, we show developmental exposure to JWH-018 reduced the locomotor activity of five dpf wild type zebrafish during dark periods in the forced light/dark test. The effects of JWH-018 were similar to the effects of THC but opposite to the effects of nicotine. The results for THC and nicotine are in line with previous studies showing a reduction in locomotion after exposure to THC [30], and an increase in locomotion after exposure to nicotine [44]. We hypothesize that cannabinoids may produce a CNS depressant effect, whereas exposure to nicotine enhances the behavioral stimulant effects of nicotine in zebrafish larvae. However, we cannot rule out that these drugs affected zebrafish behavior via impairment /activation of motor neurons or toxicity effects [45].

When anxiety-like behavior was assessed during adulthood, we observed wild-type zebrafish developmentally exposed to JWH-018 spent less time on the bottom of the tank, suggesting they were less anxious when placed in a new environment compared to non-exposed animals. These results challenge previous reports suggesting anxiogenic effects due to drug withdrawal in zebrafish [46,47]. However, none of these studies exposed fish to JWH-018, nor they exposed them at early developmental stages and tested months after withdrawal, limiting their comparability. In our study, exposure to JWH-018 started at 24 hours post fertilization, a period in which the main zebrafish brain structures (i.e. forebrain, midbrain, and hindbrain) are formed, but finer structures are still to be defined [48]. It is possible that exposures at such early ages lead to persistent adaptive changes in gene expression and neurotransmission different from the adaptive mechanisms happening during other developmental periods -such as adolescence-, which in turn may lead to different alterations in the anxiety-like responses in zebrafish in later life.

Adult zebrafish with loss-of-function mutations in *disc1* showed increased anxiety-like responses compared to wild types. These results are in line with another study showing abnormal stress response in this mutant line [49] and support the role of *disc1* in zebrafish HPI axis function [49]. Previous research in zebrafish have shown that alterations in *disc1* causes alterations in the specification of oligodendrocytes and neurons [50], and in the migration and differentiation of the neural crest (the cells that form the craniofacial cartilage and connective tissue of the head) [51]. Alterations in those processes could also underlie the alterations in behavior we observed. DISC1 is a scaffolding protein that interacts with many other proteins and regulates the formation, maintenance and correct regulation of neural networks [15]. Given the number of interacting proteins, the specific biological mechanisms by which DISC1 acts is a complex question out of the scope of this study. However, this work paves the way to using zebrafish as a legitimate model in which to investigate the role of DISC1 in stress and neurodevelopment.

We showed no evidence of *disc1* altering sensitivity to the effects of JWH-018, as the effects of JWH-018 were less appreciable in mutant zebrafish but did not reach statistical significance. These findings are in contrast with studies in mice reporting synergistic effects between THC and alterations in *Disc1*. However, disparities in the psychoactive compound (JWH-018 vs THC), in the age of exposure (early brain development vs adolescence), and in the animal model used (zebrafish vs rodents) may underlie those differences. Further work using different species is needed to replicate our findings.

There were no differences in larval behavior across *disc1* genotype groups with or without exposure to JWH-018. Interestingly, the behavioral pattern of the Tübingen wild types and the *disc1* wild type larvae in the forced light/dark test was different. Since they belonged to different zebrafish strains (Tübingen vs AB), differences may be due to their genetic background. Given the small sample sizes of the *disc1* wild type and homozygous groups (n=15-22) and the high variability in the larval behavioral responses, caution is needed before drawing strong conclusions resulting from the *disc1* larval tests as well as its comparison with the Tübingen wild types. *disc1* mutant zebrafish did not breed well: They laid less often and produced a low number of eggs, usually unfertilized. We had to perform five independent experiments and combine the results to increase the sample size, at the cost of adding experimental variation to our results. Although care was taken to ensure that time of drug exposure prior to testing, time of behavioral testing, and developmental stages were similar across experiments, these experimental parameters are known to affect zebrafish behavior [52].

JWH-018 did not affect the behavioral response of zebrafish larvae at six dpf. To maintain a gap of 48 hours between each refresh, we did not refresh the drug prior testing at this age, and therefore the absence of behavioral phenotype could be due to (1) JWH-018 metabolizes very quickly and there was no accumulation in the larvae, so after 24 hours there was no noticeable effects or (2) JWH-018 oxidates very quickly in water and its psychotropic properties were lost after a few hours in the water. In order to disentangle these scenarios, liquid chromatography-mass spectrometry analyses could be used to measure the concentrations of the drug in the water and in zebrafish tissue. It is also possible that the repeated administration of JWH-018 produced tolerance to behavioral effects in zebrafish larvae, since it has been shown that in rodents, repeated injection of similar doses of JWH-018 produced tolerance to its hypothermic and cataleptic effects [32]. Future studies where the behavioral effect of repeated vs single exposures are compared would be valuable to examine the tolerance of different drugs.

## 5. Conclusions

This is the first study looking at the behavioral effects of early developmental exposure to JWH-018 and the interaction with loss-of-function mutations in *disc1*. Our results suggest that exposure to drugs of abuse during early-development leads to long-term behavioral changes in zebrafish. However, further studies in human populations and other models are needed to confirm these findings. Our results align with previous research suggesting that functional abnormalities in DISC1 has a behavioral impact, and report no evidence of synergistic effect between developmental exposure to JWH-018 and *disc1*. These results pave the way to study molecular mechanisms by which *disc1* and developmental exposure to JWH-018 act, and give little evidence for interaction between *disc1* and developmental exposure to synthetic cannabinoids.

## Supplementary Materials

Figure S1: Response and habituation to startle stimuli test with different interstimulus intervals (ISI) in wild type zebrafish larvae, Figure S2: Tank used for novel tank diving assay.

## Author Contributions

Conceptualization, C.H.B.; methodology, C.H.B., B.D.Q., A.J.B. and J.G.G.; formal analysis, B.D.Q. A.J.B. and J.G.G.; investigation, C.H.B. and J.G.G.; resources, C.H.B.; writing—original draft preparation, C.H.B. and J.G.G.; writing—review and editing, C.H.B. and J.G.G.; visualization, J.G.G.; supervision, C.H.B.; project administration, C.H.B.; funding acquisition, C.H.B.. All authors have read and agreed to the published version of the manuscript.

## Acknowledgments

We thank Mr Luca Galantini for the assistance in animal maintenance and Miss Aida Kafai Golahmadi for preliminary work on the effects of JWH-018 on zebrafish.

## SUPPLEMENTARY MATERIAL AND APPENDICES A-B

**Figure S1.**
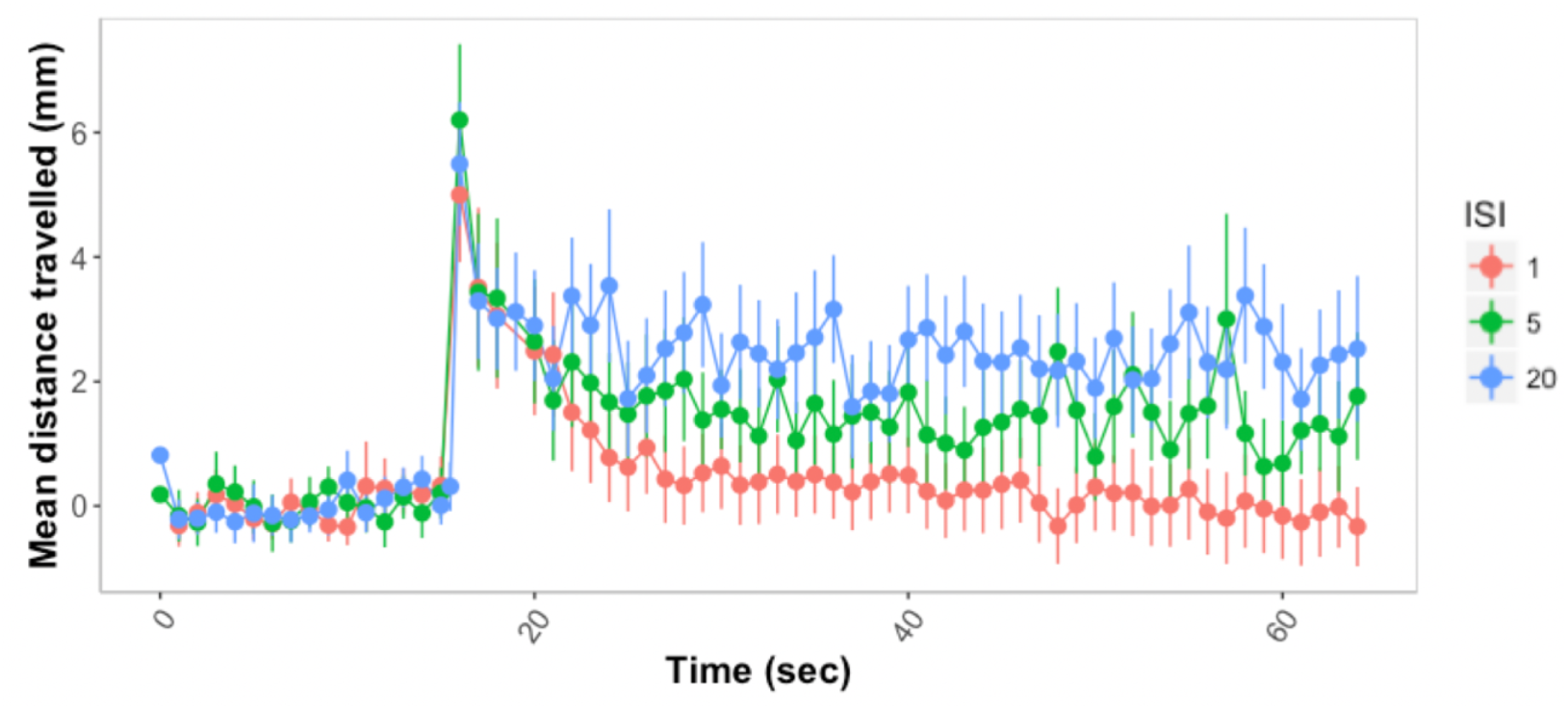
Response and habituation to startle stimuli test with different interstimulus intervals (ISI) in wild type zebrafish larvae. The first stimulus is given at second 15.

**Figure S2.**
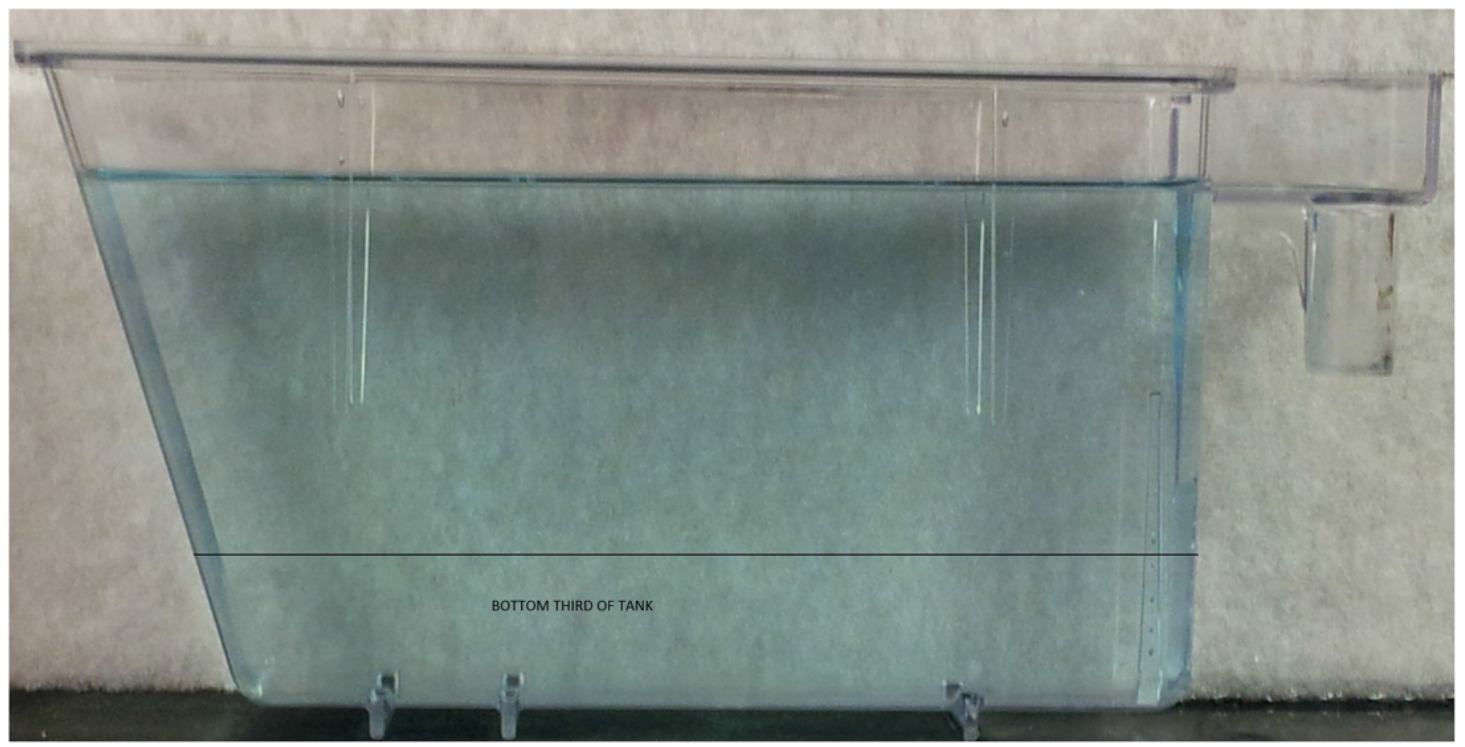
Tank used for novel tank diving assay.

## Appendix A: Developmental exposure to 0.15 μM nicotine from two to seven dpf lead to an increase in nicotine preference when fish were adults (~four months old) and conditioned to 5 μM nicotine

Drug-induced reinforcement of behavior, that reflects the hedonic value of drugs of abuse including nicotine, is highly conserved in both mammalian and non-mammalian species **[28,53–55]**. Conditioned place preference (CPP), where drug exposure is paired with specific environmental cues, is commonly used as a measure of drug-induced reinforcement and reward [56]. Previous studies have shown that zebrafish show a robust CPP to nicotine **[57–60]**.

Here, we show developmental exposure to 0.15 μM nicotine lead to altered sensitivity of the drug-induced reinforcement and reward as measured in CPP (See [57] for methodology on the CPP assay). Fish that were not developmentally treated with nicotine showed a small increase in preference when conditioned with 5 μM nicotine. By contrast, fish exposed to 0.15 μM nicotine from two to seven days showed an increased change in preference [Interaction between CPP condition and developmental exposure: F(1,73)=4.482, p=0.038] (Figure S3).

**Figure S3.**
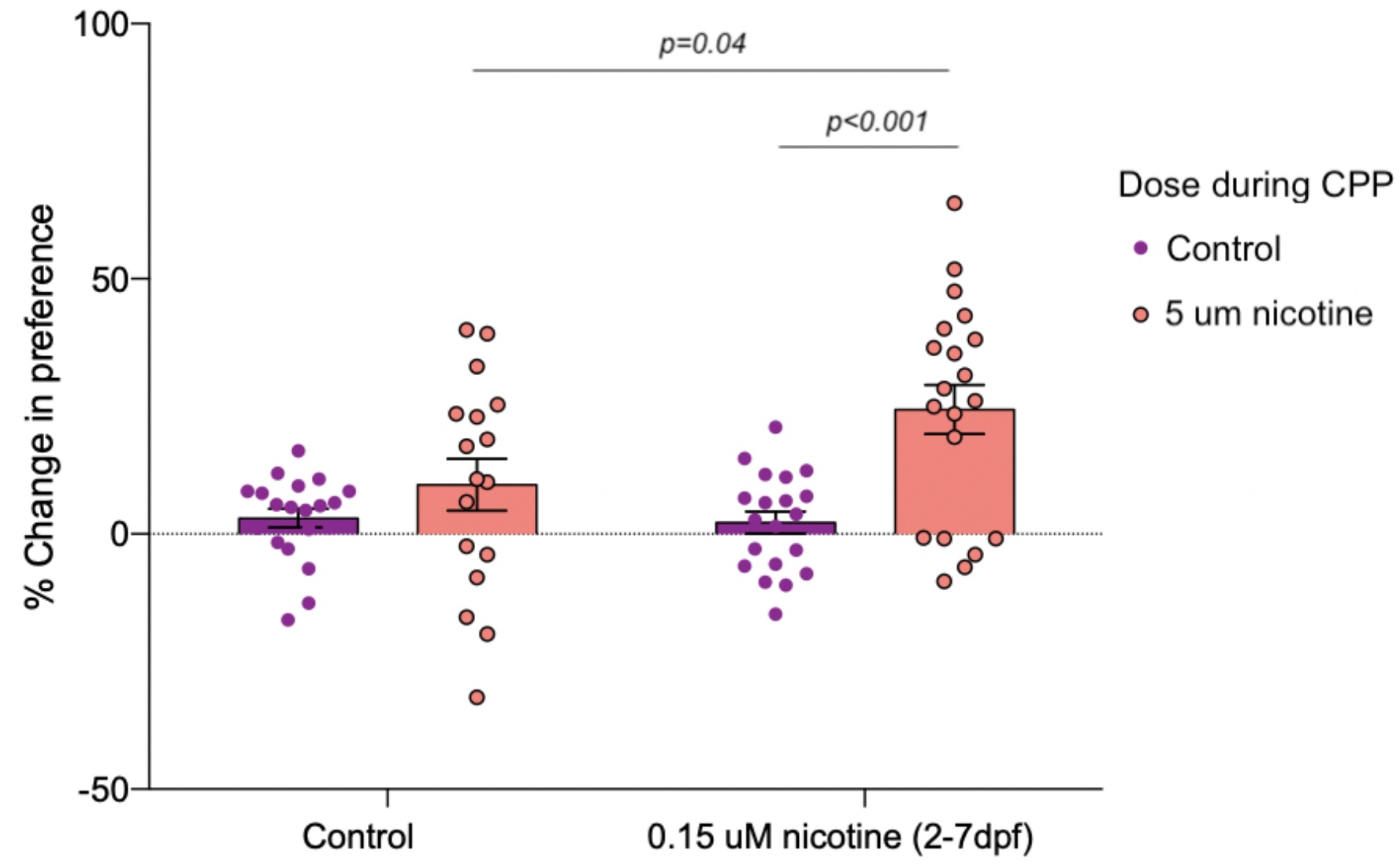
5 μM nicotine-induced place preference in adult zebrafish is exacerbated by developmental exposure to 0.15 μM nicotine (from 2-7 dpf). n=17 to 20 fish per experimental group.

## Appendix B: Behavioral assays data analysis

### Data analysis for forced/light Dark test

Firstly, we performed an overall analysis to identify the experimental variables that were significant predictors of distance travelled during the whole duration of the experiment (50 minutes). We fitted the data to a linear mixed model with total distance travelled as response variable, experimental variables (e.g. genotype, dose, time) as fixed effects, and fish ID as random effects.

We then created three subsets of the experiment: baseline, dark, and light periods. We analyzed each subset separately by fitting the data to linear mixed models as previously described. To assess differences between the first and second light periods, and between the first and second dark periods, we added the period number as fixed effect in the linear mixed models.

Linear mixed models were calculated using the R package lme4 [61]. To identify significant fixed effects, we calculated Analysis of Deviance Tables (Type II Wald χ2 tests) for the models using the R package ‘car’ [62]. Where significant differences were established, we carried out post-hoc Tukey tests with the R package ‘emmeans’ [63] to further characterize the effects.

Larvae usually increased the distance travelled during the course of the light periods. To further explore this behavior, we calculated linear models for each zebrafish at each light period using distance travelled as response variable and time as independent variable. In these linear models, the β coefficient for time represents the increase in distance travelled over time, and can be interpreted as the larva ‘recovery rate’. We constructed ANOVA models (R function ‘aov’) to assess what variables were significant predictors of the ‘recovery rate’.

### Data analysis for Habituation to startle response

We firstly investigated larvae spontaneous locomotion by testing whether distances travelled before the stimuli differed across experimental groups. We then investigated larvae startle responses by testing whether distances travelled during the stimuli differed across experimental groups. In both analyses, we fitted the data to linear mixed models using the R package lme4 [61], with total distance travelled as response variable, experimental variables (e.g. genotype, dose, time) as fixed effects, and fish ID as random effects.

### Data analysis for novel tank diving

To analyze genotype and/or treatment differences in the *time that zebrafish spent on the bottom* of the tank, we performed beta regressions using the R package ‘betareg’ [64]. We used beta regression because proportion time spent on the bottom of the tank was used as response variable. Proportion data is bounded by the interval [0, 1] and often exhibits heterogeneity in variance, which violates statistical assumptions used by linear models [64].

To analyze genotype or treatment differences in the *total distance* that zebrafish travelled in the tank, we fitted the data to a linear mixed model with the total distance travelled during one minute as response variable, time, genotype and/or treatment as fixed effects, and fish ID as random effects.

To analyze genotype or treatment differences in the *number of transitions* that zebrafish made between the top and the bottom of the tank, we fitted the data to a generalized linear mixed model with Poisson distribution. The Poisson distributions is commonly used when the response variable is count data [65]. We used the number of transitions to the top-bottom of the tank response variable, time, genotype or/and treatment as fixed effects, and fish ID as random effects.

Experiments were replicated on different days, and data was jointly analyzed afterwards. Mixed models were calculated using the R package lme4 [61]. To identify experimental variables with significant effects, we calculated Analysis of Deviance Tables (Type II Wald χ2 tests) for the models using the R package ‘car’ [62]. Where significant differences were established, we carried out post-hoc Tukey tests with the R package ‘emmeans’ [63] to further characterize the effects.

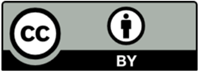 © 2020 by the authors. Submitted for possible open access publication under the terms and conditions of the Creative Commons Attribution (CC BY) license (http://creativecommons.org/licenses/by/4.0/).

